# Neural space-time model for dynamic scene recovery in multi-shot computational imaging systems

**DOI:** 10.1101/2024.01.16.575950

**Authors:** Ruiming Cao, Nikita Divekar, James Nuñez, Srigokul Upadhyayula, Laura Waller

## Abstract

Computational imaging reconstructions from multiple measurements that are captured sequentially often suffer from motion artifacts if the scene is dynamic. We propose a neural space-time model (NSTM) that jointly estimates the scene and its motion dynamics. Hence, we can both remove motion artifacts and resolve sample dynamics. We demonstrate NSTM in three computational imaging systems: differential phase contrast microscopy, 3D structured illumination microscopy, and rolling-shutter DiffuserCam. We show that NSTM can recover subcellular motion dynamics and thus reduce the misinterpretation of living systems caused by motion artifacts.

## Introduction

Multi-shot computational imaging systems capture multiple raw measurements sequentially and combine them through computational algorithms to reconstruct a final image that enhances the capabilities of the imaging system (e.g., super-resolution [8, 11], phase retrieval [20], hyperspectral imaging [14]). Each raw measurement is captured under a different condition (e.g., illumination coding, pupil coding) and hence encodes a different subset of the information. The reconstruction algorithm must then decode this information to generate the final reconstruction.

If the sample is moving during the multi-shot capture sequence, the reconstruction may be blurry or suffer artifacts [6] since the system effectively encodes information from slightly different scenes at each timepoint. Thus, most methods require that the sample be static during the full acquisition, sacrificing the system’s temporal resolution and limiting the types of samples that can be imaged. One possible approach for imaging dynamic samples is to reduce acquisition time by multiplexing measurements via hardware modifications [21, 29, 32], developing more data-efficient reconstruction algorithms [4, 9, 13], or deploying additional data priors with deep learning techniques [3, 7, 18, 23, 26, 30, 31]. However, because these methods collect less data, they come with additional trade-offs such as reduced resolution or cross-talk. They may also be impractical to implement; data priors, for example, are non-trivial to generate (e.g., due to the lack of access to the groundtruth) and may suffer issues from out-of-distribution samples [2].

Here we take another approach for imaging moving samples, where we model the sample dynamics in order to account for it during the image reconstruction. Modeling sample dynamics in multi-shot methods is challenging for two reasons: First, each measurement has a different encoding, so we cannot simply register the raw images to solve for the motion. Second, the motion can be highly complex and deformable, necessitating a pixel-level motion kernel. Deep learning offers a convenient way to develop such flexible motion models that would be very difficult to express analytically. For example, recent work successfully used a deep learning approach (with a robust data prior) to model dynamics in the case of single molecule localization microscopy [24].

We propose a neural space-time model (NSTM) that can recover a dynamic scene by modeling its spatiotemporal relationship in multi-shot imaging reconstruction. NSTM exploits the temporal redundancy of dynamic scenes. This concept, widely used in video compression, assumes that a dynamic scene evolves smoothly over adjacent timepoints. NSTM uses two coordinate-based neural networks [25, 27], one to represent the motion and the other to represent the scene, as illustrated in Fig. 1b. The motion network outputs a motion kernel for a given timepoint, which estimates the motion displacement for each pixel of the scene. Subsequently, the scene network generates a scene using spatial coordinates that have been adjusted for motion by the motion network. Then, the generated scene is passed into the multi-shot system’s forward model to produce a rendered measurement. The network weights of NSTM are optimized through gradient descent to ensure that the rendered measurements match with the acquired measurements.

**Fig. 1.**
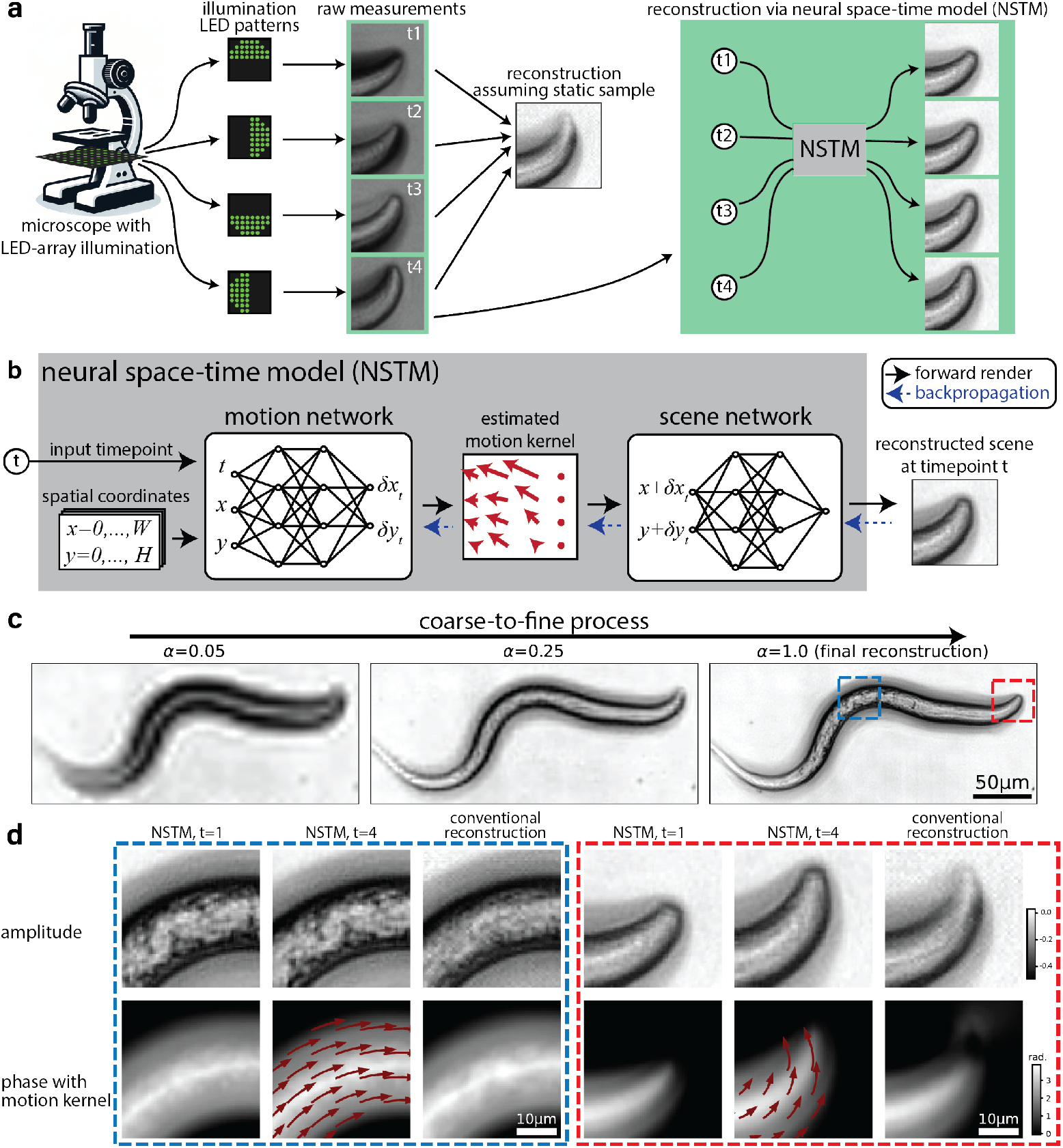
**a**, Multi-shot computational imaging systems capture a series of images under different conditions and then computationally reconstruct the final image. For example, differential phase contrast microscopy (DPC) captures four images with different illumination conditions to reconstruct quantitative phase. Sequential capture of the raw data results in motion artifacts for dynamic scenes, since the reconstruction algorithm assumes a static scene. Our proposed neural space-time model (NSTM) extends such methods to dynamic samples, by modeling and reconstructing the motion at each timepoint as well as the motion-corrected scene. **b**, NSTM consists of two coordinate-based networks, one for the motion and one for the scene. Given any timepoint as the input, NSTM generates the reconstruction at that timepoint. **c**, The coarse-to-fine process for the reconstruction of a live *C. elegans* worm imaged by DPC. **d**, Zoom-in for NSTM reconstruction at different timepoints with the recovered motion kernel overlaid, and conventional reconstruction (assuming a static scene) for comparison.

The motion and scene networks in NSTM are interdependent, and failing to synchronize their updates leads to poor convergence of the model. This poor convergence typically happens when the scene network overfits to the measurements before the motion is recovered, a situation common for scenes involving more complex motion (Extended Fig. 1 & 2). To mitigate this issue, we developed a coarse-to-fine process (detailed in Methods), which controls the granularity of the outputs from both networks. Specifically, the reconstruction starts by recovering only the low-frequency features and motion, and then gradually refines higher-frequency details and local deformable motion as illustrated in Fig. 1c.

**Fig. 2.**
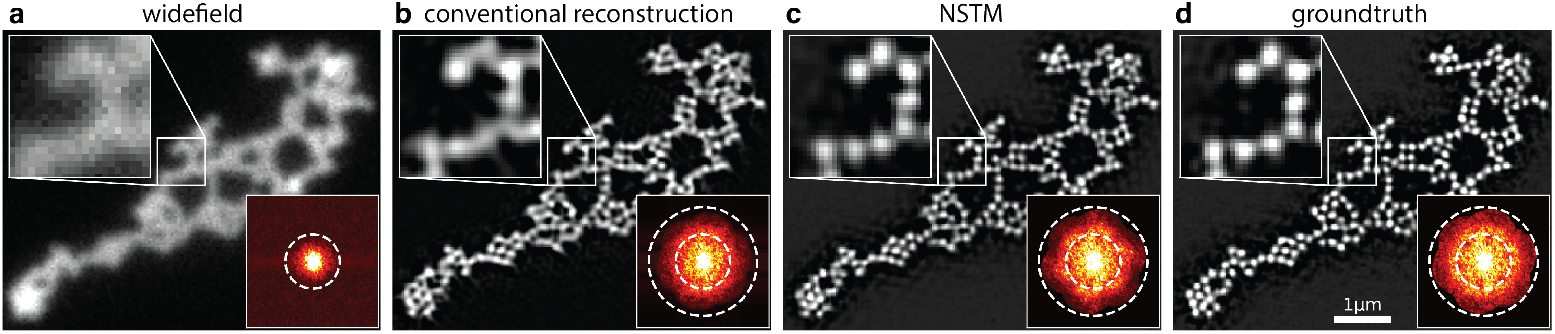
Structured illumination microscopy (SIM) super-resolved reconstruction of a dense microbead sample with vibrating motion. **a**, Diffraction-limited widefield image. **b**, The conventional SIM reconstruction algorithm (fairSIM [16]) assumes a static scene, so suffers from motion blur. **c**, Our NSTM reconstruction resolves all of the sub-resolution sized beads and gives a similar quality reconstruction as the groundtruth case (**d**), in which we collected the data without sample motion. Gamma correction with power of 0.7 is applied to all frequency spectra.

Notably, NSTM does not involve any pre-training or data priors, and as a result, it can be plugged into any multi-shot system with a differentiable and deterministic forward model. We demonstrate NSTM for three different computational imaging systems: differential phase contrast microscopy (DPC) [28], 3D structured illumination microscopy (SIM) [8] and rolling-shutter DiffuserCam [1].

## Results

### Differential phase contrast microscopy (DPC)

Our first multi-shot computational imaging system captures four raw images, and then reconstructs the amplitude and phase of a sample. The images are captured with four different illumination source patterns, which are generated by an LED array microscope in which the traditional brightfield illumination unit is replaced by a programmable LED array [22]. In Fig. 1a, we show the system and raw images captured for a live, moving *C. elegans* sample. The conventional reconstruction algorithm assumes a static scene over these four raw images. Consequently, unaccounted sample motion leads to artifacts in the reconstruction (Fig. 1d). Through the coarse-to-fine process (Fig. 1c), the NSTM recovers the motion of the *C. elegans* at each timepoint, giving a clean reconstruction without motion artifacts.

### 3D structured illumination microscopy (SIM)

Our second multi-shot system is 3D SIM [8] which captures 15 raw measurements at each *z* plane (three illumination orientations, five phase shifts for each orientation). The conventional 3D SIM reconstruction assumes there is no motion during the acquisition; thus, it is limited to fixed samples. Previous work in extending 3D SIM to live cells focuses on accelerating the acquisition through faster hardware [5, 15, 32] or assumes translation-only motion [8]. NSTM provides a strategy to recover and account for deformable motion from raw images captured by 3D SIM. Because we model motion during the acquisition of a single volume, we can reconstruct both the super-resolved image and the dynamics (see Methods).

Figure 2a shows results for a single-layer dense microbead sample in which we introduced motion by gently pushing and releasing the optical table during the acquisition. Using a conventional reconstruction algorithm (fairSIM [16]) results in a low-resolution image in which the individual beads cannot be resolved due to motion blur (Fig. 2b). In contrast, as shown in Fig. 2c, our NSTM reconstruction recovers the motion (Extended Fig. 3b and d) and resolves individual beads with a quality comparable to the groundtruth reconstruction. In this case, the groundtruth was reconstructed from a separate set of raw measurements captured without motion (Fig. 2d).

Applying this technique to live-cell imaging, Fig. 3 and Extended Fig. 4 show 3D SIM reconstructions for a live RPE-1 cell expressing StayGold-tagged mitochondrial matrix protein. In Fig. 3b, the conventional reconstruction appears to show a mitochondrion with a tubule branch (red arrow); however, our NSTM result recovers the sample dynamics (see Extended Fig. 4b and Extended Movie 3) and thus recognizes that it is actually a single tubule which is moving during the acquisition time (Fig. 3d). This can be further verified by the low-resolution widefield images (Fig. 3e) and by running our NSTM algorithm without the motion update (Extended Fig. 4c). In addition to resolving motion, NSTM removes motion blur, recovering features that were blurred in the conventional reconstruction (blue arrows in Fig. 3b-c), and thus NSTM preserves more high-frequency content compared with conventional reconstructions (Extended Fig. 4d).

**Fig. 3.**
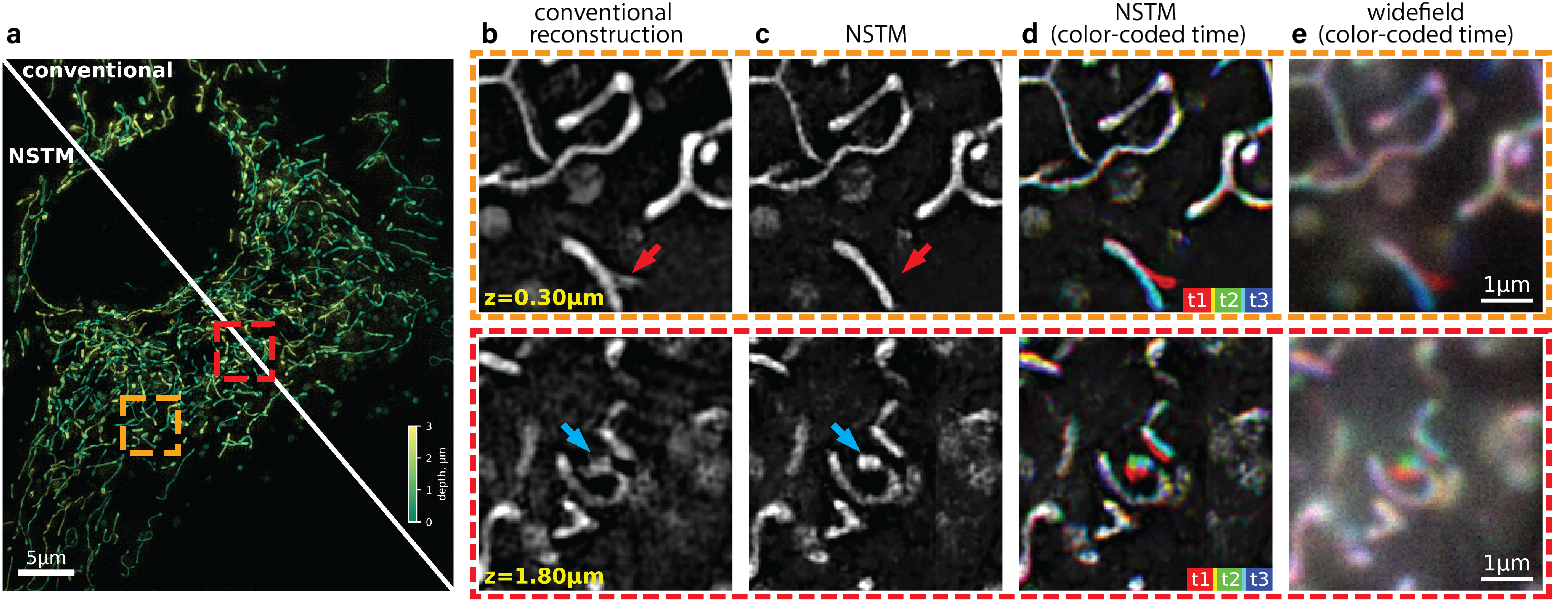
3D SIM reconstruction of a live RPE-1 cell expressing StayGold-tagged mitochondrial matrix protein. **a**, Maximum projection of the volume with color-coded depth. **b-c**, Zoom-in single-plane reconstruction comparison between conventional 3D SIM reconstruction algorithm (CUDA-accelerated three-beam SIM reconstruction software [8]) and our NSTM algorithm. **d-e**, The NSTM reconstructions and widefield images at three timepoints coded by colors. The widefield images are obtained by summing the raw images from five phase shifts.

**Fig. 4.**
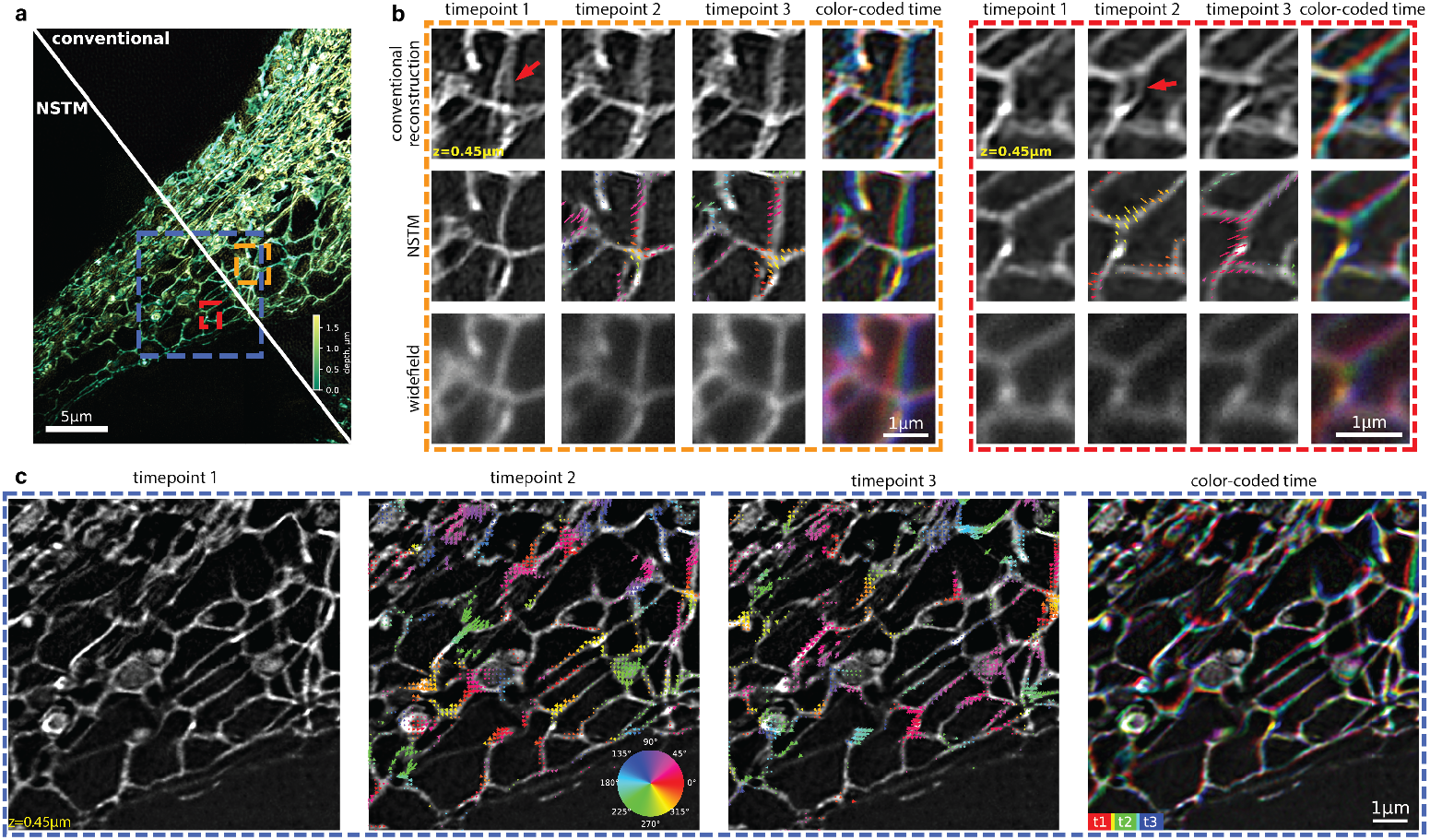
3D SIM experimental results for a RPE-1 cell expressing StayGold-tagged endoplasmic reticulum (ER). **a** Maximum projection of the reconstructed volume with color-coded depth. **b**, Zoom-in single-plane reconstructions and widefield images at three timepoints, each of which corresponds to a different illumination orientation. Moving window approach (see Methods) is used for the conventional reconstruction (CUDA-accelerated three-beam SIM reconstruction software [8]) at different timepoints. The NSTM reconstructions are overlaid with the recovered motion kernels which show the sample’s motion displacements from the previous timepoint. The colors of the motion kernel indicate motion directions. **c**, Single-plane zoom-in NSTM reconstructions at three timepoints and the combined view with color-coded time. The motion kernels on the second and third timepoints show the structure’s motion displacements from the previous timepoint, with the color-coding to indicate motion directions.

In another experiment, we imaged a live RPE-1 cell expressing StayGold-tagged endoplasmic reticulum (ER) using 3D SIM (Fig. 4). The conventional reconstruction struggles to resolve clear ER network structures, likely due to their fast dynamics (see red arrows). Additionally, the motion artifacts in the conventional reconstruction often have distinct appearances at different timepoints (Fig. 4b), making it difficult to interpret the ER dynamics from a time-series reconstruction. NSTM, on the other hand, recovers the motion kernels and the dynamic scene from the same set of raw images for a single volume reconstruction, and the ER structures it resolves are consistent over time. The recovered motion kernels reveal the dynamics happening at different timepoints within a single 3D SIM acquisition as shown in Fig. 4c and Extended Movie 4.

### Rolling-shutter DiffuserCam lensless imaging

Our third multi-shot computational imaging example is rolling-shutter DiffuserCam [1], a lensless camera that compressively encodes a high-speed video into a single captured image. This method leverages the fact that each row of the image, captured sequentially by the rolling shutter, contains information about the whole scene at that timepoint, due to the system’s large point-spread-function (PSF) generated by a optical diffuser. To enable this video reconstruction from the single raw image, the original algorithm [1] uses total variation (TV) regularization to promote smoothness. In contrast, by modeling for the motion explicitly, NSTM produces cleaner reconstructions without over-smoothing (Extended Fig. 6b). As a byproduct of NSTM, the motion trajectory for any point can be queried directly from the motion network (Extended Fig. 6c).

## Discussion

We demonstrated our neural space-time model (NSTM) for recovering motion dynamics and removing motion-induced artifacts in three different multi-shot imaging systems; however, the models are general and should find use in other multi-shot computational imaging methods. Notably, NSTM does not use any data priors or pre-training. Hence, it is compatible with any multi-shot system with a differentiable and deterministic forward model. For multi-shot imaging systems like 3D SIM, which do not use gradient-based reconstruction, we can alternatively implement a forward model as part of the NSTM reconstruction.

One limitation of our method is that it hinges on the assumption of temporal redundancy, implying moderate overall motion across measurements and correlatable scenes at adjacent timepoints. Despite that the two-network design of NSTM allows an explicit motion model and ensures reconstruction fidelity, it also introduces an additional constraint: since the scene network does not depend on the temporal coordinate, any frame of a dynamic scene has to be obtained by deforming a static reconstruction (from the scene network) with a motion kernel (from the motion network). These two assumptions limit the NSTM’s applicability to fluctuating scenes (such as neuron firing or fluorescence photoactivation) or scenes with appearing/disappearing imaging features. To overcome this limit, future work could modify the NSTM architecture to account for the fluctuation and/or incorporate the time-dependency to the scene network.

Another limitation is that our NSTM reconstructions generally require more computation than conventional methods. For example, the dense microbead reconstruction using NSTM took about 10 minutes on a Nvidia Titan V GPU, in contrast to the conventional algorithm (fairSIM) which completed in less than 10 seconds on a CPU, and the live cell 3D reconstruction using NSTM took 3 hours on eight Nvidia A100 GPUs. Future work could improve the computational efficiency of NSTM by better initialization of network weights, hyper-parameter search for a faster convergence, and data-driven methods to optimize a part of the model in a single pass.

One interesting advantage of NSTM is that it uses coordinate-based neural networks to effectively store and represent both the scene and its motion dynamics. Unlike convolutional neural networks (CNNs) which always require a matrix as the input, a coordinate-based network can accommodate arbitrary values whose coordinates may not be on a matrix grid. This is especially advantageous for modeling spatiotemporal relationships, as it can handle sub-pixel motion shifts and non-uniformly sampled measurements, without requiring interpolation for a uniformly sampled matrix. A potential extension for NSTM is the frame interpolation for a high frame-rate reconstruction, which can be achieved by oversampling timepoints during the reconstruction rendering, as demonstrated in Extended Movie 5.

In summary, we showed that our NSTM method can recover motion dynamics and thus resolve motion artifacts, in multi-shot computational imaging systems, using only the typical datasets used for conventional reconstructions. The ability to recover dynamic samples within a single multi-shot acquisition seems particularly promising for observing subcellular systems in live biological samples. By accounting for motion through NSTM’s joint reconstruction, NSTM reduces the risk of misinterpretations in the study of living systems caused by motion artifacts in multi-shot acquisitions. Further, it effectively increases the temporal resolution of the system when multi-shot data is captured.

## Methods

### Sample preparation

The dense microbead sample was made with 0.19μm dyed microbeads (Bangs Laboratories, FC02F). The stock solution was diluted 1:100 with distilled water and placed on a glass-bottom 35mm dish coated by Poly-L-lysine solution (Sigma Aldrich, P8920).

The RPE-1 cell lines were cultured using Dulbecco’s Modified Eagle Medium/Nutrient Mixture F-12 (Thermo Scientific 11320033) supplemented with 10% FBS (VWR Life Science 100% Mexico Origin 156B19), 2mM L-Glutamine, 100 Units/mL penicillin, and 100mg/mL streptomycin (Fisher Scientific 10378016). Trypsin-EDTA (0.25%) phenol red (Fisher Scientific 25200114) was used to detach cells for passaging. To generate the cell lines, we obtained the pCSII-EF/mt-(n1)StayGold (Addgene plasmid #185823) and pcDNA3/er-(n2)oxStayGold(c4)v2.0 (Addgene plasmid #186296) from Atsushi Miyawaki [10] to tag the mitochondrial matrix and the endoplasmic reticulum, respectively. The er-(n2)oxStayGold(cr)v2.0 sequence was PCR amplified and cloned into a lentiviral vector containing an EF1 alpha promoter. The vector is a derivative of Addgene #60955 with the sgRNA sequence removed. Lentiviral particles containing each plasmid were produced by transfecting standard packaging vectors along with the plasmids into HEK293T cells using TransIT-LT1 Transfection Reagent (Mirus, MIR2306). Media was changed at 24 hours post-transfection, without disturbing the adhered cells, and the viral supernatant was harvested approximately 50 hours post transfection. The supernatant was filtered through 0.45 mm PVDF syringe filter and about 1 mL was used to directly seed a 10 cm plate of hTERT RPE-1 cells (ATCC CRL-4000). Two days post-infection, cells were analyzed on BD FACSAria Fusion Sorter and BDFACSDiva Software and the highest 5% of StayGold/GFP fluorescence cells were sorted for each line (gating strategy illustrated in Extended Fig. 7) and expanded for imaging experiments.

### Data acquisition

The 3D SIM images were acquired on a commercial three-beam SIM system (Zeiss Elyra PS.1) using an oil immersion objective (Zeiss, 100*×* 1.46 NA) and 1.6x tube lens. The effective pixel size is 40.6nm. The system captures 15 images at each plane, with 3 illumination orientations and 5 phase shifts for each orientation. A single image plane was acquired for the dense microbead sample. 20 planes with a step size of 150nm were captured for the RPE-1 cell expressing StayGold-tagged mitochondrial matrix protein, and 12 planes with a step size of 150nm were imaged for the RPE-1 cell expressing StayGold-tagged endoplasmic reticulum. The SIM system has a illumination update delay of around 20ms for each phase shift or *z* -position shift, and a delay of 300ms for each illumination orientation change. We set the exposure time to 20ms for the dense microbeads and 5ms for the both cell lines.

The DPC images were obtained from [12] using a commercial inverted microscope (Nikon TE300) with 10*×* 0.25NA objective (Nikon) and an effective pixel size of 0.454μm. A LED-array [22] (SCI Microscopy) was attached to the microscope in place of the Köhler illumination unit. Four half circular illumination patterns, with the maximum illumination NA equal to the objective NA, were sequentially displayed on the LED array to capture four raw images as in [28]. Exposure time was 25ms.

The rolling shutter DiffuserCam image was obtained from its original work [1]. The raw image was taken by a color sCMOS (PCO Edge 5.5) in slow-scan rolling shutter mode (27.52μs readout time for each row) with dual shutter readout and 1320μs exposure time. The acquisition of the raw image took 31.0ms.

### NSTM implementation

The motion and the scene network of NSTM are both coordinate-based neural networks [25, 27], multi-layer perceptrons that learn a mapping from coordinates to signals. To enhance the capacity and efficiency of a coordinate-based network, we use hash embedding [17] to store multiple grids of features at different resolutions and transform a coordinate vector to a multi-resolution feature vector, ***f*** = [*f*_0_, *f*_2_,*⃛, f*_*N*−1_], before passing it into the network. As the input coordinate varies, a fine resolution feature (e.g., *f*_*N*−1_) changes more rapidly than a coarse resolution feature (e.g., *f*_0_). During the coarse-to-fine process, we re-weight the output features of the hash embedding using a granularity value, *α*, to control the granularity of the network. *α* is set by the ratio of the current epoch to the end epoch of the coarse-to-fine process, which is set to 80% of the total number of reconstruction epochs in practice. As in [19], each feature *f*_*i*_ is weighted by 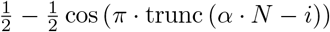, where trunc truncates a value to [0, 1]. In this way, finer features will be weighted to 0 until *α* gets larger, as illustrated in Fig. 1c.

In the forward process of NSTM (Fig. 1b), every spatial coordinate of the scene, (***x***), is concatenated with the temporal coordinate *t* and fed into the motion network. The motion network produces the motion displacement vector, *δ****x*** for each input spatiotemporal coordinate, and (***x*** + *δ****x***) is then fed into the scene network for the reconstruction value at (***x***, *t*). This process is repeated for all spatial coordinates to obtain the reconstructed scene at *t*. The scene network outputs a single channel as the fluorescent density for 3D SIM, two channels as the amplitude and phase for DPC, and three channels as RGB intensity for DiffuserCam.

To update the network weights of NSTM, the reconstructed scene is passed into the imaging system’s forward model for a rendered measurement. By comparing the rendered measurement with the actual measurement acquired in the experiment, we compute the mean square error (MSE) loss and minimize it by back-propagating its gradient to update the network weights. In our implementation, the motion network has two hidden layers with a width of 32, and the scene network has two hidden layers with a width of 128. For the conventional reconstruction of NSTM without motion update (in Extended Fig. 3a and Extended Fig. 4c), we keep all settings the same as the NSTM reconstruction except that the motion network is not updated and the input timepoints are set to zero.

### DPC reconstruction

The raw images of DPC are normalized by the background intensity. We use the linear transfer functions derived in [28] as the forward model. The conventional reconstruction is obtained by solving a Tikhonov regularization with a regularization weight of 10^−4^ for both amplitude and phase terms [28]. For ease of comparison, we add the same Tikhonov regularization to the loss term for NSTM reconstruction.

### 3D SIM reconstruction

The conventional 3D SIM reconstruction uses five measurements of different sinusoidal phase shifts to separate the complex spectra of three frequency bands and then shifts each band accordingly based on its corresponding modulation frequency. The band separation process necessitates the assumption of a static scene over those five measurements. To avoid this static assumption and preserve the temporal information, we implement the 3D SIM forward model in real space without band separation, rendering each measurement independently from NSTM’s reconstruction at the timepoint that the actual measurement is taken.

In our comparisons, we use the same illumination parameters estimated from measurements [8, 15] for both conventional reconstruction algorithms and NSTM. For the conventional reconstructions shown in Fig. 4b, we use the moving window approach to select a set of raw images around a certain timepoint to feed into the reconstruction algorithm, and we repeat this process to get the conventional reconstruction at every illumination orientation. For example, the conventional reconstruction at timepoint 3 in Fig. 4b uses raw images from illumination orientation 2 & 3 from the current acquisition^1^ and also the illumination orientation 1 from the next acquisition, where there is no delay between two acquisitions.

### Rolling-shutter DiffuserCam reconstruction

Each row of the raw image captured by rolling-shutter DiffuserCam is the time integral of the dynamic scene convolved by the caustic point-spread-function (PSF) over the rolling shutter exposure. Thus, its forward model can be written in a discrete-time sum of *T* timepoints [1], such as the raw image 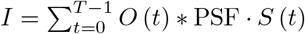, where *O* is the dynamic scene, *S* is a binary indicator of the shutter on/off state, and ∗ denotes convolution operation. However, rendering the entire image at once requires obtaining NSTM’s reconstructed scenes at all timepoints, which will be intensive on GPU memory. To make this feasible on common GPUs, during each step of the reconstruction we render a subset of image rows by only obtaining the reconstructed scenes at timepoints which have contributed signal to these rows, such that *I* (*y*) = Σ_*t* s.t.*S*(*y,t*)=1_*O* (*t*) ∗ PSF *· S* (*t*).

## Funding Acknowledgments

This work was supported by Weill Neurohub Investigators Program, CZI grant DAF2021-225666 and grant DOI https://doi.org/10.37921/192752jrgbnh from the Chan Zuckerberg Initiative DAF, an advised fund of Silicon Valley Community Foundation (funder DOI 10.13039/100014989), and STROBE: A National Science Foundation Science & Technology Center under Grant No. DMR 1548924 (NSF Grant 1351896). Laura Waller is a Chan Zuckerberg Biohub investigator. Research reported in this publication was supported in part by the National Institutes of Health S10 program under award number 1S10OD018136-01. The content is solely the responsibility of the authors and does not necessarily represent the official views of the National Institutes of Health.

## Extended Data

**Extended Movie 1**: Simulation of differential phase contrast microscopy (DPC) with various types of motion.

**Extended Movie 2**: Simulation of 3D structured illumination microscopy (SIM) with various types of motion.

**Extended Movie 3**: The 3D rendering of NSTM reconstruction for a live RPE-1 cell expressing StayGold-tagged mitochondrial matrix protein.

**Extended Movie 4**: The raw images and reconstructions for a RPE-1 cell expressing StayGold-tagged endoplasmic reticulum (ER) on a series of 3D SIM acquisitions. The recovered motion kernel with color-coded motion direction is plotted on NSTM reconstruction.

**Extended Movie 5**: DPC reconstruction of *C. elegans* with time oversampling during the rendering.

**Extended Fig. 1.**
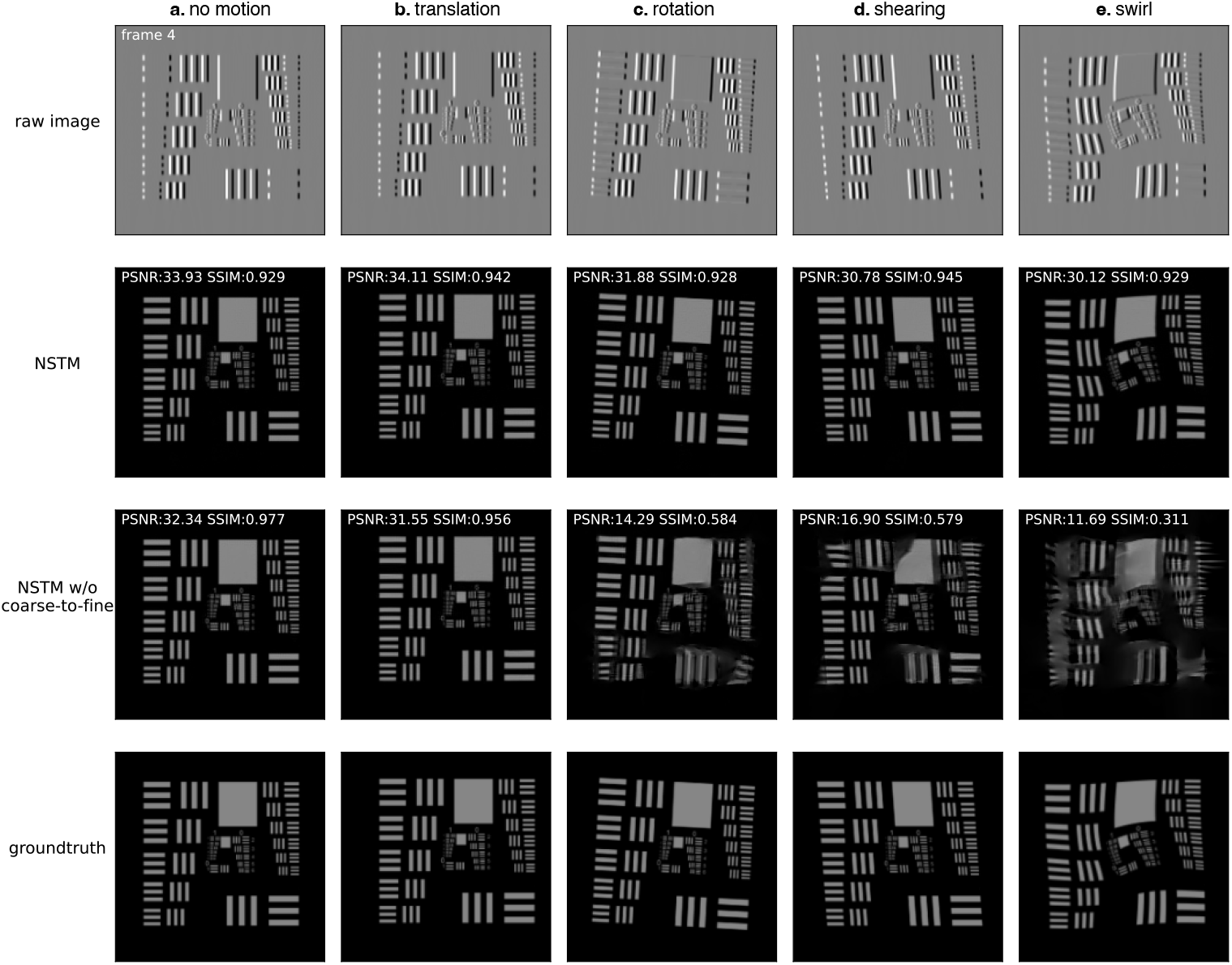
Simulations of NSTM reconstructions with various types of motion: **a**, no motion, **b**, rigid motion - translation, **c**, rigid motion - rotation, **d**, non-rigid global motion - shearing, and **e**, local deformable motion - swirl. Here, the data is simulated differential phase contrast microscopy (DPC) images of a phase-only USAF-1951 resolution target. Two reconstruction quality metrics are calculated: peak signal-to-noise ratio (PSNR) and the structural similarity index measure (SSIM). The NSTM does well with all types of motion. However, without using our coarse-to-fine process (‘NSTM w/o coarse-to-fine’), it is likely to fail as the motion gets complicated, due to poor convergence of the joint optimization of motion and scene. Full videos of the dynamic reconstruction can been seen in Extended Movie 1.

**Extended Fig. 2.**
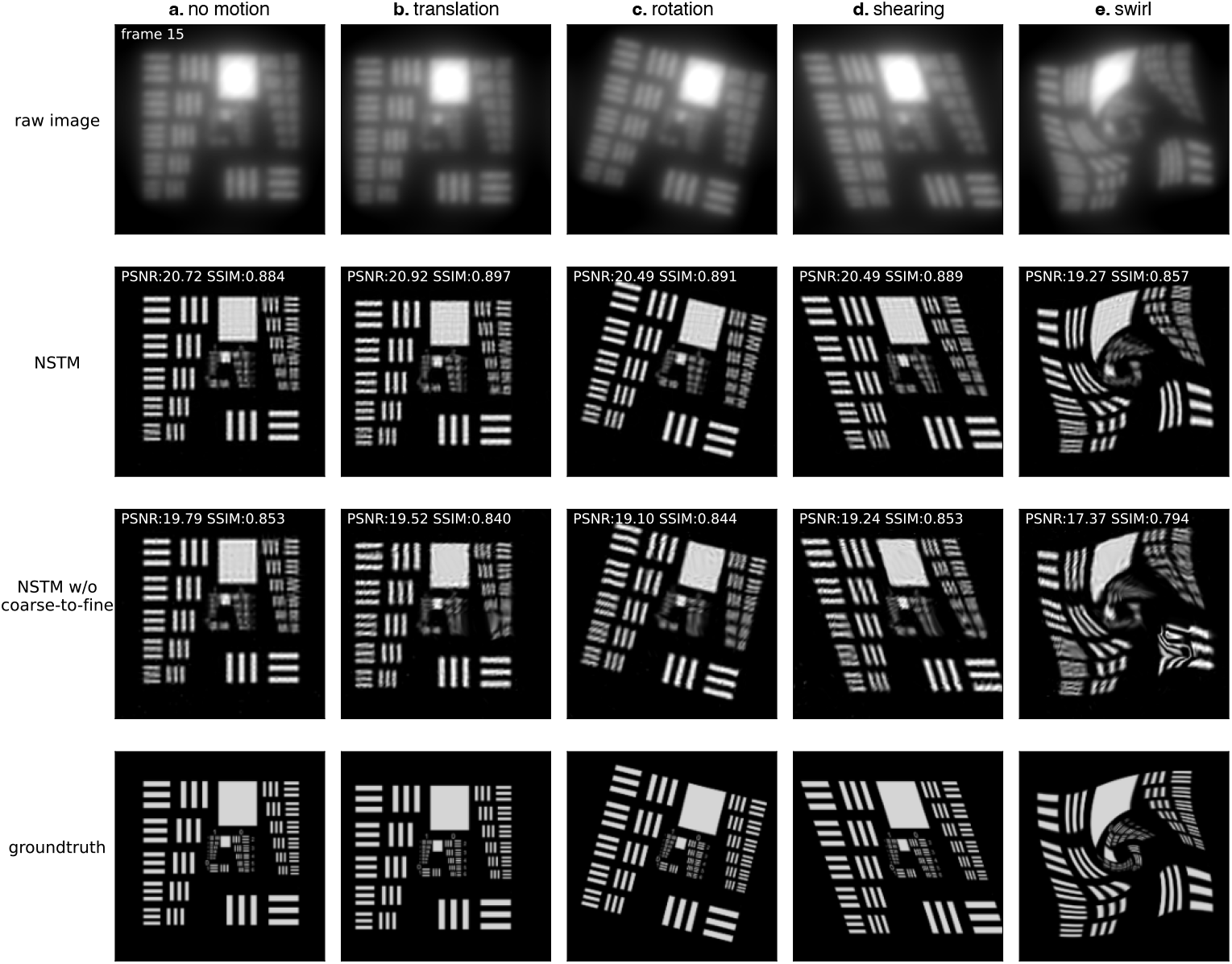
Simulations of 3D structured illumination microscopy (SIM) using fluorescent USAF-1951 resolution target with various types of motion: **a**, no motion, **b**, rigid motion - translation, **c**, rigid motion - rotation, **d**, non-rigid global motion - shearing, and **e**, local deformable motion - swirl. Full videos of the dynamic reconstruction can been seen in Extended Movie 2.

**Extended Fig. 3.**
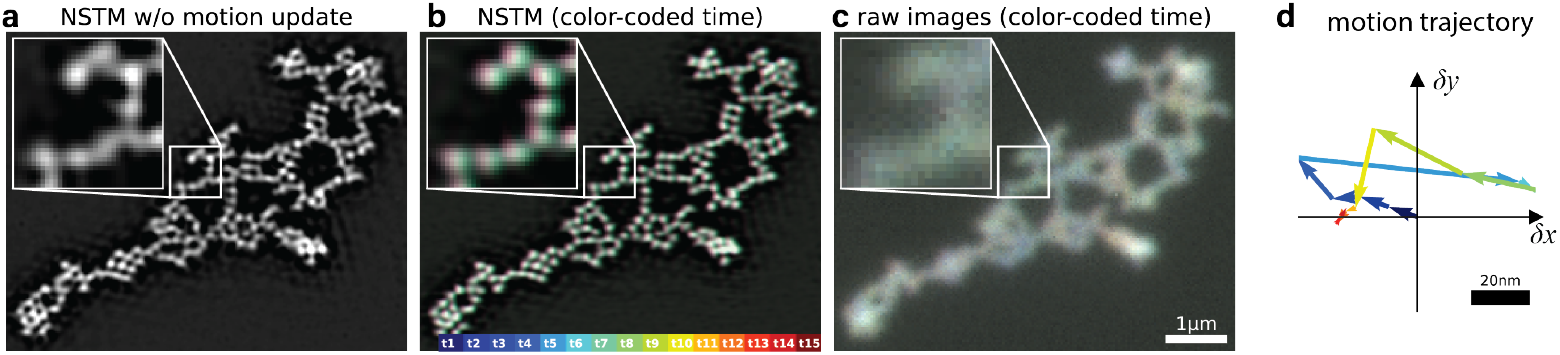
Additional results on the dense microbead sample from Fig. 2: **a**, Reconstruction using NSTM without the motion update results in motion blurring similar to the conventional reconstruction in Fig. 2b, since dynamics are not accounted for. **b**, NSTM reconstruction with color-coded time. **c**, The raw images with color-coded time. In the images with color-coded time, each timepoint of raw images or reconstruction is drawn in a distinct color as indicated by the color bar. The ‘color dispersion’ in the zoom-in reconstruction suggests that a subtle motion is recovered by NSTM. **d**, The recovered motion trajectory of a pixel on the vibrating microbeads from NSTM reconstruction. Each arrow shows the motion displacement vector with respect to the previous timepoint as indicated by the color code.

**Extended Fig. 4.**
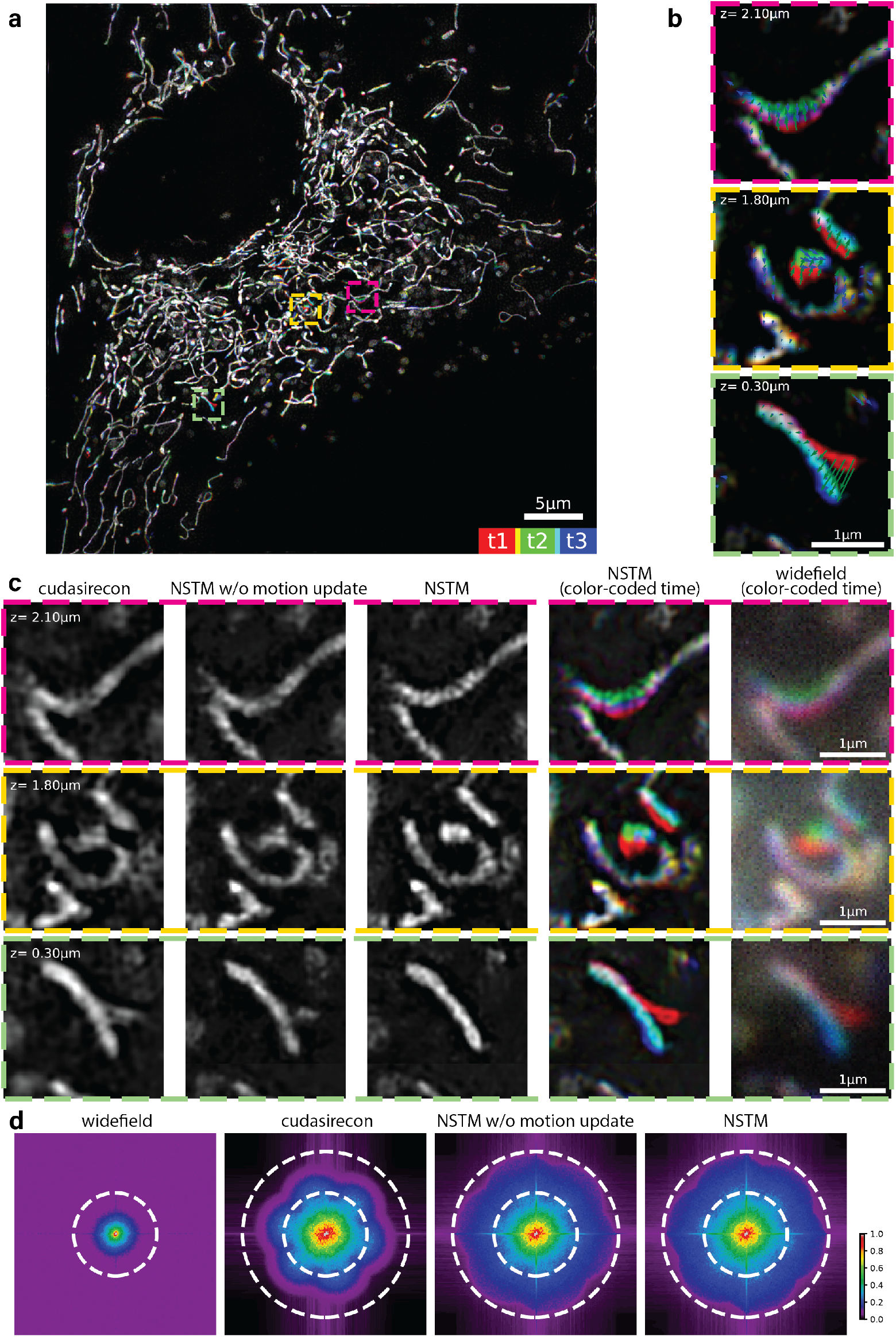
Additional 3D SIM results for the mitochondria-labeled RPE-1 cell from Fig. 3. **a**, Maximum projection of NSTM reconstruction volume, with three colors denoting the three time-points that correspond to the three illumination orientations. **b**, Single-plane zoom-in of the NSTM reconstruction with color-coded time. The overlaid vector fields show the motion displacement recovered by NSTM, with their colors to indicate their corresponding timepoints. **c**, Single-plane zoom-in comparisons, from left to right: conventional reconstructions (CUDA-accelerated three-beam SIM reconstruction software (‘cudasirecon’) [8] and NSTM without motion update), NSTM reconstruction, NSTM reconstruction with color-coded time (three colors for three illumination orientations), and widefield images with color-coded time. **d**, A comparison of their spatial frequency spectra. The two dotted circles indicate the diffraction-limited bandwidth and SIM super-resolved bandwidth respectively. Gamma correction with power of 0.5 is applied to all frequency spectra for better contrast.

**Extended Fig. 5.**
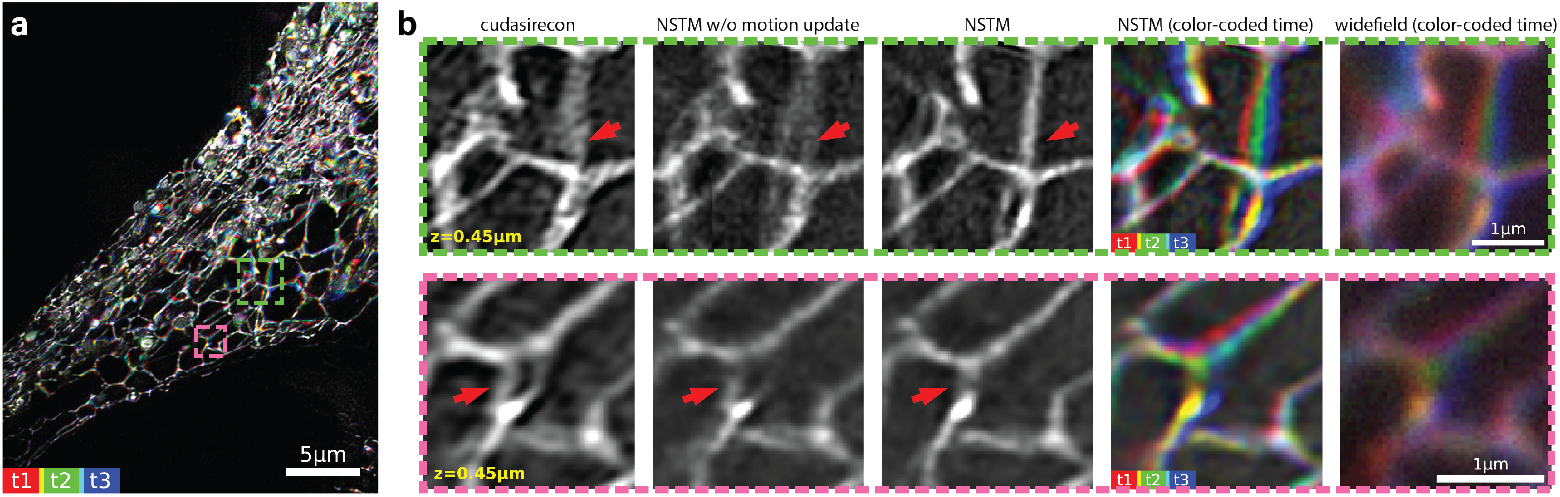
Additional 3D SIM results for the live endoplasmic reticulum-labeled RPE-1 cell. **a**, Maximum z-projection of NSTM reconstruction volume, with three colors denoting the three time-points that correspond to the three illumination orientations. **b**, Single-plane zoom-in comparisons, from left to right: conventional reconstructions (CUDA-accelerated three-beam SIM reconstruction software (‘cudasirecon’) [8] and NSTM without motion update), NSTM reconstruction, NSTM reconstruction with color-coded time (three colors for three illumination orientations), and widefield images with color-coded time.

**Extended Fig. 6.**
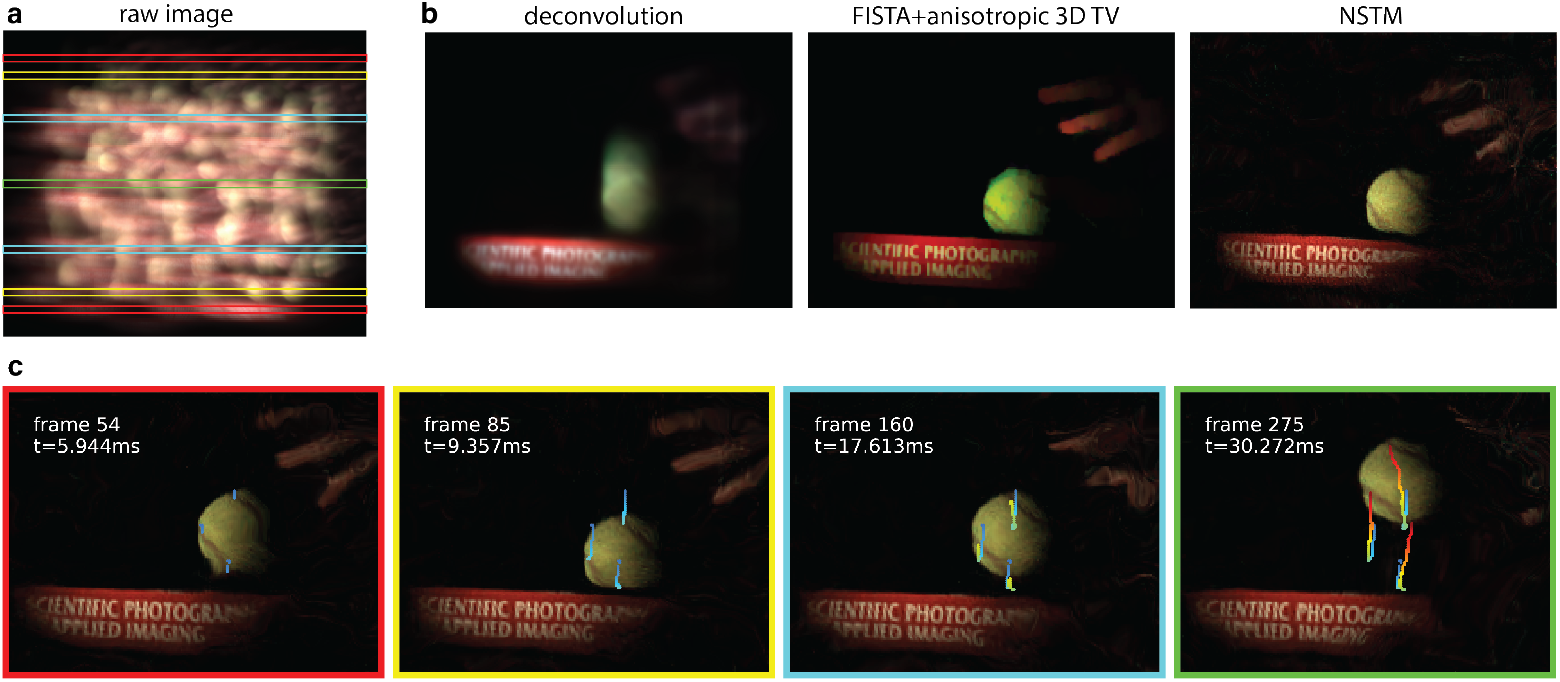
Results for rolling-shutter DiffuserCam. **a**, The raw image measurement. **b**, Comparisons of the reconstruction using basic deconvolution (assumes a static scene), FISTA with anisotropic 3D Total Variation regularization (TV) [1] (the original reconstruction method), and our NSTM algorithm. **c**, NSTM reconstruction at different timepoints, with their corresponding measurement rows indicated by colored boxes on the raw image. The colored curves show some selected motion trajectories recovered by the motion network.

**Extended Fig. 7.**
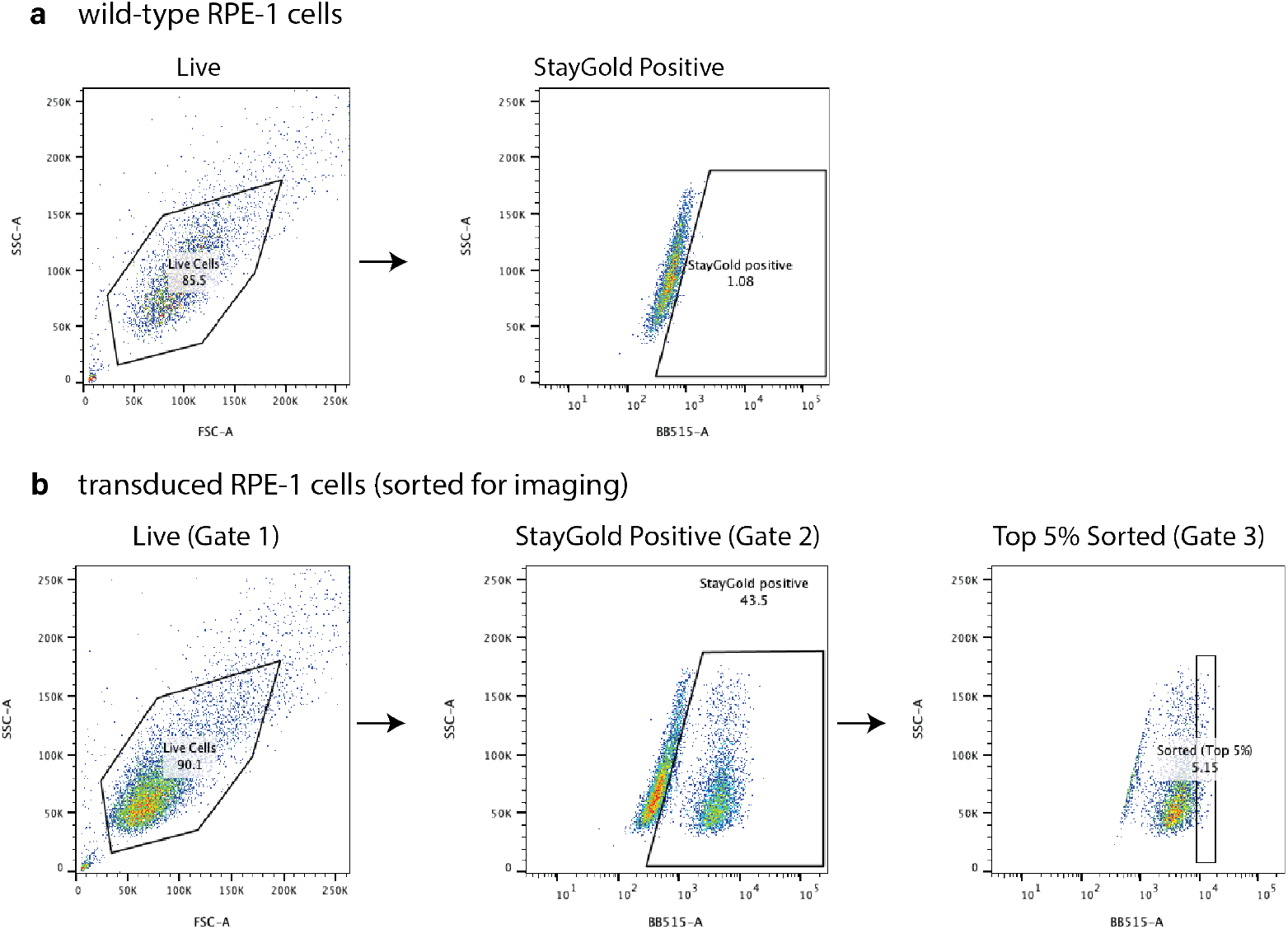
Gating strategy for cell sorting. **a**, Wild-type RPE-1 cells were used to gate for Live Cells (Gate 1) and the StayGold negative cells were used to gate for the StayGold positive population (Gate 2). **b**, To sort samples that were transduced with StayGold expressing plasmids, Gate 1 (Live cells) was applied followed by Gate 2 (StayGold positive), and then top 5% of the StayGold positive cell population (Gate 3) was sorted using BDFACS Aria Fusion Sorter and expanded using DMEM-F12 for subsequent imaging experiments.

## Footnotes

1 We use the term ‘acquisition’ to refer to ‘timepoint’ in a regular context of time-series acquisition, since ‘timepoint’ is already heavily used for time within a single acquisition of a scene in this manuscript.

## References

[1] Antipa N, Oare P, Bostan E, et al (2019) Video from stills: Lensless imaging with rolling shutter. In: International Conference on Computational Photography, IEEE, pp 1–8

[2] Belthangady C, Royer LA (2019) Applications, promises, and pitfalls of deep learning for fluorescence image reconstruction. Nature methods 16(12):1215–1225

[3] von Chamier L, Laine RF, Jukkala J, et al (2021) Democratising deep learning for microscopy with zerocostdl4mic. Nature communications 12(1):2276

[4] Dertinger T, Colyer R, Iyer G, et al (2009) Fast, background-free, 3d superresolution optical fluctuation imaging (sofi). Proceedings of the National Academy of Sciences 106(52):22287–22292

[5] Fiolka R, Shao L, Rego EH, et al (2012) Time-lapse two-color 3d imaging of live cells with doubled resolution using structured illumination. Proceedings of the National Academy of Sciences 109(14):5311–5315

[6] Förster R, Wicker K, Müller W, et al (2016) Motion artefact detection in structured illumination microscopy for live cell imaging. Optics Express 24(19):22121–22134

[7] Ge B, He Y, Deng M, et al (2022) Single-frame label-free cell tomography at speed of more than 10,000 volumes per second. arXiv preprint arXiv:220203627

[8] Gustafsson MG, Shao L, Carlton PM, et al (2008) Three-dimensional resolution doubling in wide-field fluorescence microscopy by structured illumination. Biophysical journal 94(12):4957–4970

[9] Gustafsson N, Culley S, Ashdown G, et al (2016) Fast live-cell conventional fluorophore nanoscopy with imagej through super-resolution radial fluctuations. Nature communications 7(1):12471

[10] Hirano M, Ando R, Shimozono S, et al (2022) A highly photostable and bright green fluorescent protein. Nature Biotechnology 40(7):1132–1142

[11] Huang B, Bates M, Zhuang X (2009) Super-resolution fluorescence microscopy. Annual review of biochemistry 78:993–1016

[12] Kellman M, Chen M, Phillips ZF, et al (2018) Motion-resolved quantitative phase imaging. Biomedical Optics Express 9(11):5456–5466

[13] Laine RF, Heil HS, Coelho S, et al (2023) High-fidelity 3d live-cell nanoscopy through data-driven enhanced super-resolution radial fluctuation. Nature Methods pp 1–8

[14] Lu G, Fei B (2014) Medical hyperspectral imaging: a review. Journal of biomedical optics 19(1):010901–010901

[15] Lu-Walther HW, Kielhorn M, Förster R, et al (2015) fastsim: a practical implementation of fast structured illumination microscopy. Methods and Applications in Fluorescence 3(1):014001

[16] Müller M, Mönkemöller V, Hennig S, et al (2016) Open-source image reconstruction of super-resolution structured illumination microscopy data in imagej. Nature communications 7(1):10980

[17] Müller T, Evans A, Schied C, et al (2022) Instant neural graphics primitives with a multiresolution hash encoding. ACM Transactions on Graphics (ToG) 41(4):1–15

[18] Nehme E, Weiss LE, Michaeli T, et al (2018) Deep-storm: super-resolution singlemolecule microscopy by deep learning. Optica 5(4):458–464

[19] Park K, Sinha U, Barron JT, et al (2021) Nerfies: Deformable neural radiance fields. In: Proceedings of the IEEE/CVF International Conference on Computer Vision, pp 5865–5874

[20] Park Y, Depeursinge C, Popescu G (2018) Quantitative phase imaging in biomedicine. Nature photonics 12(10):578–589

[21] Phillips ZF, Chen M, Waller L (2017) Single-shot quantitative phase microscopy with color-multiplexed differential phase contrast (cdpc). PloS one 12(2):e0171228

[22] Phillips ZF, Eckert R, Waller L (2017) Quasi-dome: A self-calibrated high-na led illuminator for fourier ptychography. In: Imaging Systems and Applications, Optica Publishing Group, pp IW4E–5

[23] Qiao C, Li D, Guo Y, et al (2021) Evaluation and development of deep neural networks for image super-resolution in optical microscopy. Nature Methods 18(2):194–202

[24] Saguy A, Alalouf O, Opatovski N, et al (2023) Dblink: Dynamic localization microscopy in super spatiotemporal resolution via deep learning. Nature Methods pp 1–10

[25] Sitzmann V, Martel J, Bergman A, et al (2020) Implicit neural representations with periodic activation functions. Advances in neural information processing systems 33:7462–7473

[26] Speiser A, Müller LR, Hoess P, et al (2021) Deep learning enables fast and dense single-molecule localization with high accuracy. Nature methods 18(9):1082–1090

[27] Stanley KO (2007) Compositional pattern producing networks: A novel abstraction of development. Genetic programming and evolvable machines 8:131–162

[28] Tian L, Waller L (2015) Quantitative differential phase contrast imaging in an led array microscope. Optics express 23(9):11394–11403

[29] Waller L, Kou SS, Sheppard CJ, et al (2010) Phase from chromatic aberrations. Optics express 18(22):22817–22825

[30] Weigert M, Schmidt U, Boothe T, et al (2018) Content-aware image restoration: pushing the limits of fluorescence microscopy. Nature methods 15(12):1090–1097

[31] Wu Y, Rivenson Y, Wang H, et al (2019) Three-dimensional virtual refocusing of fluorescence microscopy images using deep learning. Nature methods 16(12):1323–1331

[32] York AG, Chandris P, Nogare DD, et al (2013) Instant super-resolution imaging in live cells and embryos via analog image processing. Nature methods 10(11):1122–1126

